# Metabolomics of lung microdissections reveals region- and sex-specific metabolic effects of acute naphthalene exposure in mice

**DOI:** 10.1101/2021.06.08.447459

**Authors:** Nathanial C. Stevens, Patricia C. Edwards, Lisa M. Tran, Xinxin Ding, Laura S. Van Winkle, Oliver Fiehn

## Abstract

Naphthalene is a ubiquitous environmental contaminant produced by combustion of fossil fuels and is a primary constituent of both mainstream and side stream tobacco smoke. Naphthalene elicits region-specific toxicity in airway club cells through cytochrome P450 (P450)-mediated bioactivation, resulting in depletion of glutathione and subsequent cytotoxicity. While effects of naphthalene in mice have been extensively studied, few experiments have characterized global metabolomic changes in the lung. In individual lung regions, we found metabolomic changes in microdissected mouse lung conducting airways and parenchyma obtained from animals sacrificed 2, 6, and 24 hours following naphthalene treatment. Data on 577 unique identified metabolites were acquired by accurate mass spectrometry-based assays focusing on lipidomics and non-targeted metabolomics of hydrophilic compounds. Statistical analyses revealed distinct metabolite profiles between the two major lung regions. In addition, the number and magnitude of statistically significant exposure-induced changes in metabolite abundance were different between lung airways and parenchyma for unsaturated lysophosphatidylcholines (LPCs), dipeptides, purines, pyrimidines, and amino acids. Importantly, temporal changes were found to be highly distinct for male and female mice, with males exhibiting predominant treatment-specific changes only at two hours post-exposure. In females, metabolomic changes persisted until six hours post-naphthalene treatment, which may explain the previously characterized higher susceptibility of female mice to naphthalene toxicity. In both males and females, treatment-specific changes corresponding to lung remodeling, oxidative stress response, and DNA damage were observed, which may provide insights into potential mechanisms contributing to the previously reported effects of naphthalene exposure in the lung.

## Introduction

Naphthalene is a ubiquitous polycyclic aromatic hydrocarbon emitted into the atmosphere by combustion of fossil fuels, cigarette smoke, biomass burning, and several other sources (1). Humans are exposed to naphthalene primarily through inhalation but can also ingest naphthalene through diet (2). Widespread human exposure to naphthalene is of concern due to findings from animal studies that demonstrate acute toxicity as well as formation of neoplasms in rodents, prompting the classification of naphthalene as a potential human carcinogen (3). The proposed mechanism of naphthalene toxicity is through cytochrome P450 (P450) monooxygenase mediated bioactivation. CYP2F2, the predominant isoform responsible for metabolizing naphthalene in the mouse lung, rapidly metabolizes naphthalene into a reactive epoxide (4). This epoxide is detoxified via conjugation with glutathione, but can form DNA and protein adducts upon glutathione depletion, which is accompanied by cytotoxicity in the airway epithelium (5, 6). The physiological effects of naphthalene exposure include apical membrane blebbing and oxidative stress followed by changes in energy supply and ultimately loss of cells in the airway epithelium (7). The human ortholog, CYP2F1, has much lower activity towards naphthalene relative to CYP2F2, which may suggest that humans are at a lower risk for naphthalene-induced tumor formation (8, 9). However, transgenic expression of CYP2F1 and CYP2A13, another P450 isoform expressed in human lungs, was sufficient to bioactivate inhaled naphthalene in vivo and mediate naphthalene’s respiratory toxicity in humanized mice (10).

Non-ciliated lung airway epithelial cells, commonly referred to as club cells, highly express CYP2F2 and are highly susceptible to naphthalene induced injury in mice (11, 12). Club cells are most abundant in the distal airways of mice and in the respiratory bronchioles of non-human primates and humans (11). Club cell expression of CYP2F2 in mice is related to site-specific toxicity following naphthalene exposure. Additionally, female mice tend to be more susceptible to the toxic effects of naphthalene, highlighting the importance of both target region- and sex-specific effects of exposure (13). Most metabolomics studies use whole organs. However, in the case of naphthalene, the cellular targets of toxicity are club cells which are confined to the conducting airways. Nonetheless, lung regions that are not targeted for toxicity and that contain distinct cell types such as the alveolar cell types found in lung parenchyma may contribute to the initial response. These studies are needed to better understand the mechanisms of naphthalene toxicity that could lead to adverse outcomes in the lung in both target and non-target regions for acute cytotoxicity.

Metabolomics enables global characterization of metabolites produced by an organism and metabolic changes associated with toxicant exposure or environmental interactions (14, 15). Previous studies have implemented nontargeted metabolomics analyses to characterize changes in metabolism in response to naphthalene, demonstrating significant alterations with respect to treatment (16, 17). However, these analyses have been limited to the sampling of homogenized whole lung lobes, precluding the ability to distinguish metabolic responses between different lung regions. Identifying these region-specific responses is especially important for toxicants that target specific cell types with heterogenous distribution throughout the lung (18). One potential technique to isolate lung regions is gross lung microdissection, which has previously been established as an approach to distinguish differences in response to naphthalene exposure between lung airways and the surrounding parenchyma (19).

Our objective was to characterize metabolic responses to naphthalene in microdissected lung tissue from male and female mice using nontargeted metabolomics. We treated male and female C57BL/6 mice with a single i.p. injection of naphthalene and sampled gross microdissected airways and parenchyma at 2, 6, and 24 hours post-injection. Liquid chromatography - accurate mass tandem mass spectrometry (LC-MS/MS) assays for both lipids and hydrophilic metabolites were implemented to maximize coverage of annotations for both types of tissues. A series of multivariate and univariate statistical analyses was performed to identify metabolite changes among various groups. We hypothesized that metabolite profiles would differ both between tissue types and between treatments. Based on previous studies, we also anticipated female metabolite profiles to be perturbed more than males in response to treatment.

## Materials and Methods

### Animal protocol

Adult male and female C57BL/6 mice (Envigo, Inc.) aged 8-10 weeks were housed on a 12/12 light/dark cycle and fed a diet consisting of Purina 5001 lab diet. Each animal received an i.p. injection of either corn oil, which was used as a vehicle control, or naphthalene dissolved in corn oil (200 mg/kg); all mice were treated at the same time of day, in the morning. Mice were euthanized at 2, 6, or 24 hours post-injection with a lethal injection of pentobarbital and necropsied immediately following euthanasia. Lungs from each mouse were cannulated, removed en bloc, and inflated using a heated solution of 1% agarose (w/v) in 0.01M phosphate buffered saline (PBS). The left lobe of each mouse was microdissected following a previous protocol (19). The resulting airways and parenchyma were immediately stored at −80°C until analysis. All animal experiments were conducted under approved protocols reviewed by the UC Davis Institutional Animal Care and Use Committee in accordance with guidelines for animal research established by the National Institutes of Health.

### Preparation of samples and LC/MS/MS analysis

Frozen microdissected tissues were lyophilized for approximately 24 hours. Dried samples were homogenized, and 1 mg of dried tissue was used for analysis, roughly equivalent to 10 mg of fresh tissue. Tissue homogenates were extracted on ice in 2-mL centrifuge tubes by adding 225 μL of methanol and an internal standard mixture included in **Table E1** and 750 μL of methyl tert-butyl ether containing cholesterol ester 22:1 (20). The top and bottom fractions were evaporated to dryness, which contained hydrophobic and hydrophilic metabolites, respectively. The dry samples containing hydrophilic metabolites were resuspended in 110 μL of 80% acetonitrile, 20% water, and an internal standard mixture included in **Table E2.** The dry samples containing hydrophobic metabolites were resuspended in 100 μL of 90% methanol, 10% toluene, and 50 ng/ml CUDA. Detailed methods for extraction and resuspension can be found in the online data supplement. All samples were analyzed by a ThermoFisher Scientific Vanquish UHPLC+ liquid chromatography system coupled to a Q-Exactive HF orbital ion trap mass spectrometer. Detailed analysis, instrument, and chromatography parameters are included the online data supplement.

### Data processing and Statistics

Data processing was completed in MS-DIAL v.4.18 (22). Identification for all compounds was based on mass spectra from *in silico* libraries, MassBank of North America (https://massbank.us), and NIST20. Experimental spectra from MassBank of North America are publicly available and NIST20 spectra are commercially available for use. Matches were determined based on m/z, retention time, and MS/MS fragmentation pattern (23). The processed data were normalized using Systematic Error Removal in Random Forest (SERRF) (24), a machine-learning algorithm that normalizes experimental samples based on systematic variation in pooled QC samples. Statistical analysis was completed in R v.3.6.1 on the log-transformed dataset. One-way ANOVA was performed with Tukey’s HSD post-hoc test to adjust for multiple comparisons. Multivariate statistical analysis was conducted by principal component analysis (PCA), hierarchical clustering analysis (HCA) and chemical similarity enrichment analysis (ChemRICH) (25). Volcano plots and heatmaps were generated in R using the Bioconductor packages EnhancedVolcano and ComplexHeatmap (26, 27).

## Results

### Metabolomic and lipidomics compound annotations

A total of 577 unique metabolites were annotated in the dataset across both lipophilic and hydrophilic chromatographic platforms and both electrospray modes. Unknown chromatographic features within the dataset were excluded from the final analysis. The annotations between lipidomics and HILIC analysis were broken down into major classes and is shown in **Figure 1**. A full list of compounds and classes for all platforms is included online in a supplemental datasheet. Among the annotated metabolites analyzed in the study, the values for median relative standard deviation of pooled experimental samples used as a measure of technical variance were 8.3% and 15.5% for compounds identified by lipidomics and HILIC, respectively. Most lipids identified in the dataset were neutral lipids, with a slightly lower number of phospholipids identified. Each major class of lipid was also categorized based on the degree of unsaturation, which can be attributed to biological function. The largest class of hydrophilic metabolites was annotated as derivatives of amino acids, included dipeptides.

**Figure 1.**
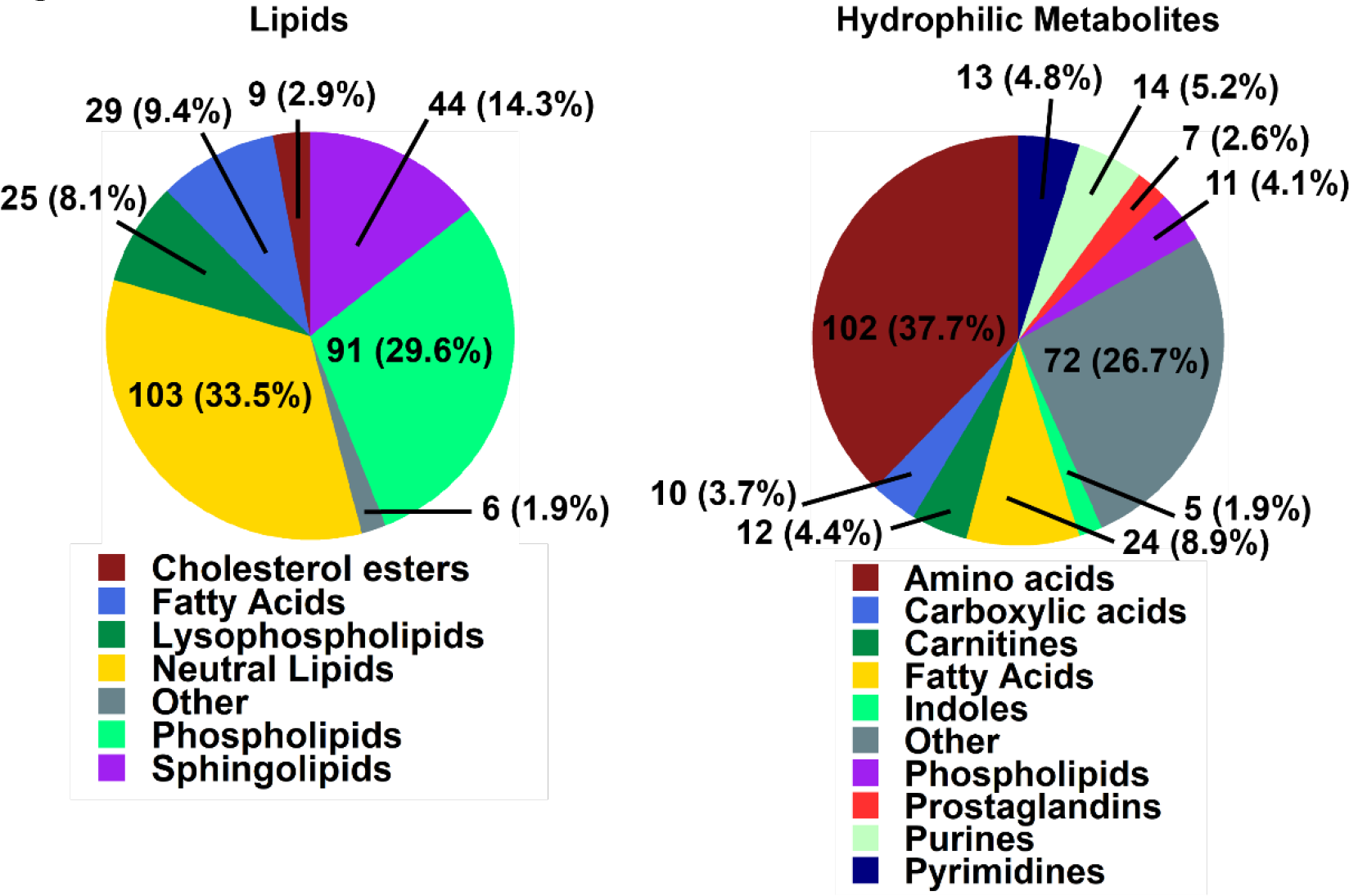
Overview of annotations identified by lipidomics and HILIC analysis. *Left*: complex lipids were classified by ClassyFire software into seven major lipid classes, consisting of 307 unique annotations. *Right*: Hydrophilic compounds were classified into 10 major metabolite classes, comprising 270 unique annotations. ClassyFire categories with less than 5 compounds were summarized into “Other” class labels.

### Metabolic differences between mouse airways and parenchyma dominate overall variance

We first determined which experimental factors contributed most to overall differences in the data set, using principal components analysis (PCA) as an unbiased multivariate dimension reduction technique (**Figure 2**). Principal component 1 (PC1) accounted for almost 20% of the overall data variance that was likely due to biological variation between sexes, treatments, and timepoints. Technical errors did not contribute to PC1 variance as seen by the close clustering of the quality control pool samples (**Figure 2**). No single sample needed to be removed due to potential gross difference to all other samples. The next vector, PC2, explained 14.3% of the total data variance, sufficient to completely distinguish lung airways and parenchyma samples within the data set. However, differences attributed to sex, naphthalene exposure or temporal changes did not dominate metabolic phenotypes to an extent that would lead to overt clustering along axes of PCA plots. Instead, these biological differences led to overall variance with slowly decreasing importance, leading to only 50% explained variance combined by the top-5 principal components (**Figure E1**).

**Figure 2.**
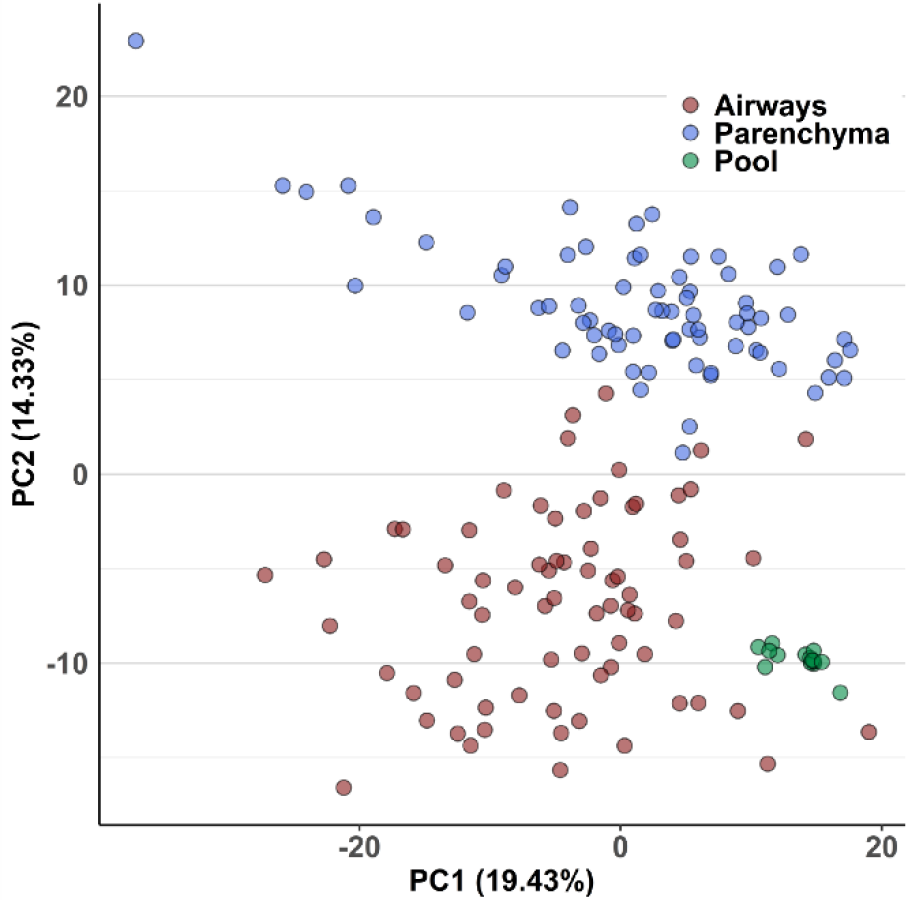
Principal Components Analysis (PCA) of metabolic variance in mouse lungs. PC2 discriminates metabolic phenotypes of mouse airways and parenchyma. Pool samples were prepared by mixing fractions of each extracted parenchyma and airways sample, which were used as a measure of technical variance of the analytical method.

### Statistical analysis of metabolite profiles between tissues and naphthalene treatment

Multivariate analysis identifying changes in metabolite classes between treatments and between tissues was next used to determine differences that contributed to the observed variation by PCA. Chemical enrichment similarity analysis (ChemRICH) enabled characterization of significantly altered metabolite classes in response to naphthalene treatment for both lung airways and parenchyma. ChemRICH is a multivariate statistical approach used as an alternative to traditional pathway mapping that does not rely on database size and groups each metabolite based on its chemical structure, which often alludes to a compound’s biological function(25). Initial ChemRICH analyses between naphthalene-treated and control animals revealed distinct differences at each timepoint and between sexes. In males, amino acids, purines, and several other metabolite classes were altered in response to treatment at 2 hours post-injection (**Figure E2**). However, no significant changes in metabolite classes were identified by ChemRICH at 6 or 24 hours in males. In contrast, changes in metabolite classes in females were present both at 2 hours (**Figure E3**) and at 6 hours, with the most extensive changes identified at 6 hours post-injection. Dipeptides and unsaturated LPCs were decreased in both airways and parenchyma following naphthalene treatment at 6 hours in females, whereas amino acid species were both increased and decreased following treatment (**Figure 3A-B**). Additionally, multiple pyrimidine nucleosides were decreased in parenchyma but not in airways of the naphthalene-treated animals, highlighting the difference in response between the two tissues (**Figure 3B**). For significantly altered metabolite classes in both airways and parenchyma of females, the average metabolite abundance of each class yielded the greatest fold difference at 6 hours comparing the two treatment groups (**Figure 3C-D**). Lastly, ChemRICH analysis comparing metabolite classes between tissues for each sex revealed striking differences in the lipid profiles of airways and parenchyma samples, which were dominated by substantially higher levels of triacylglycerides (TG) in airways than in parenchyma. Importantly, these differences did not appear to be mediated by treatment or sex, as the difference in TG abundance was evident at each timepoint in the control-treated male and female mice (**Figure E4**).

**Figure 3.**
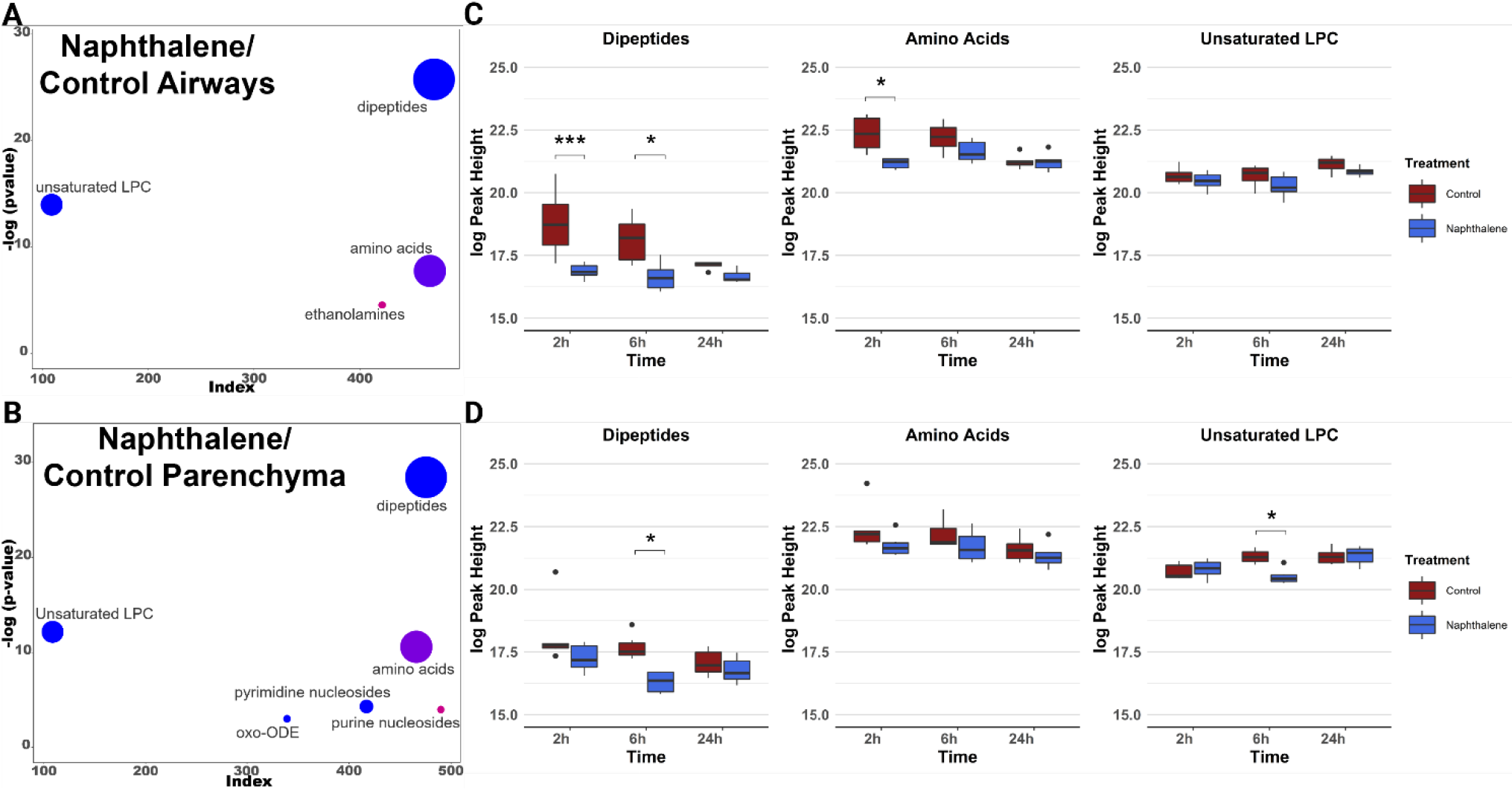
Metabolite profiles of lung airways and parenchyma are altered in response to naphthalene-treatment in females 6 hours post-injection. **A-B)** ChemRICH plots comparing naphthalene-treated airways and parenchyma in female mice 6 hours post-injection, respectively. The size of each circle represents the relative number of metabolites contained within each cluster. Red circles indicate all metabolites increase within a cluster, while blue circles indicate all metabolites decrease within a cluster. Pink and purple circles represent a mix consisting of mostly increased and decreased metabolite abundances, respectively. Axes correspond to the −logP value of a metabolite class plotted against index values assigned to each metabolite in the online datasheet included as supplemental material. P-values used for the input of each ChemRICH were calculated by one-way ANOVA with Tukey’s post-hoc analysis. P-values for each ChemRICH cluster were calculated using the Kolmogorov-Smirnov test. **C-D)** Boxplots displaying the average intensities for the largest clusters of metabolite classes altered in female airways and parenchyma for all timepoints, respectively. Axes represent the log_10_ peak height of each sample for each timepoint, and samples with values greater than 1.5 times the interquartile range are indicated by dots on each plot. * p<0.05, *** p<0.001.

Following the analysis of metabolite class changes, we next wanted to evaluate alterations within each significantly altered class to identify underlying changes in subclasses of metabolites related to a specific biological function. Due to the absence of significant differences in metabolite classes in male mice after 2 hours, we focused our subsequent analyses on female mice tissues sampled 6 hours post-injection. Hierarchical clustering analysis (HCA) comparing between both tissues and treatments demonstrated several significantly altered metabolites of the same subclass. Once again, unsaturated TGs were more abundant in airways than in parenchyma in the control mice, which were all clustered following HCA. Two clusters, one consisting of unsaturated cholesterol esters (CE) and another consisting of unsaturated phosphatidylcholines (PC) and LPCs were all lower in abundance in airways than in parenchyma in control-treated mice. Interestingly, the fold change in LPCs and PCs decreased in response to naphthalene treatment, whereas the fold change in TGs between both tissues was greatly increased following treatment (**Figure 4A**).

**Figure 4.**
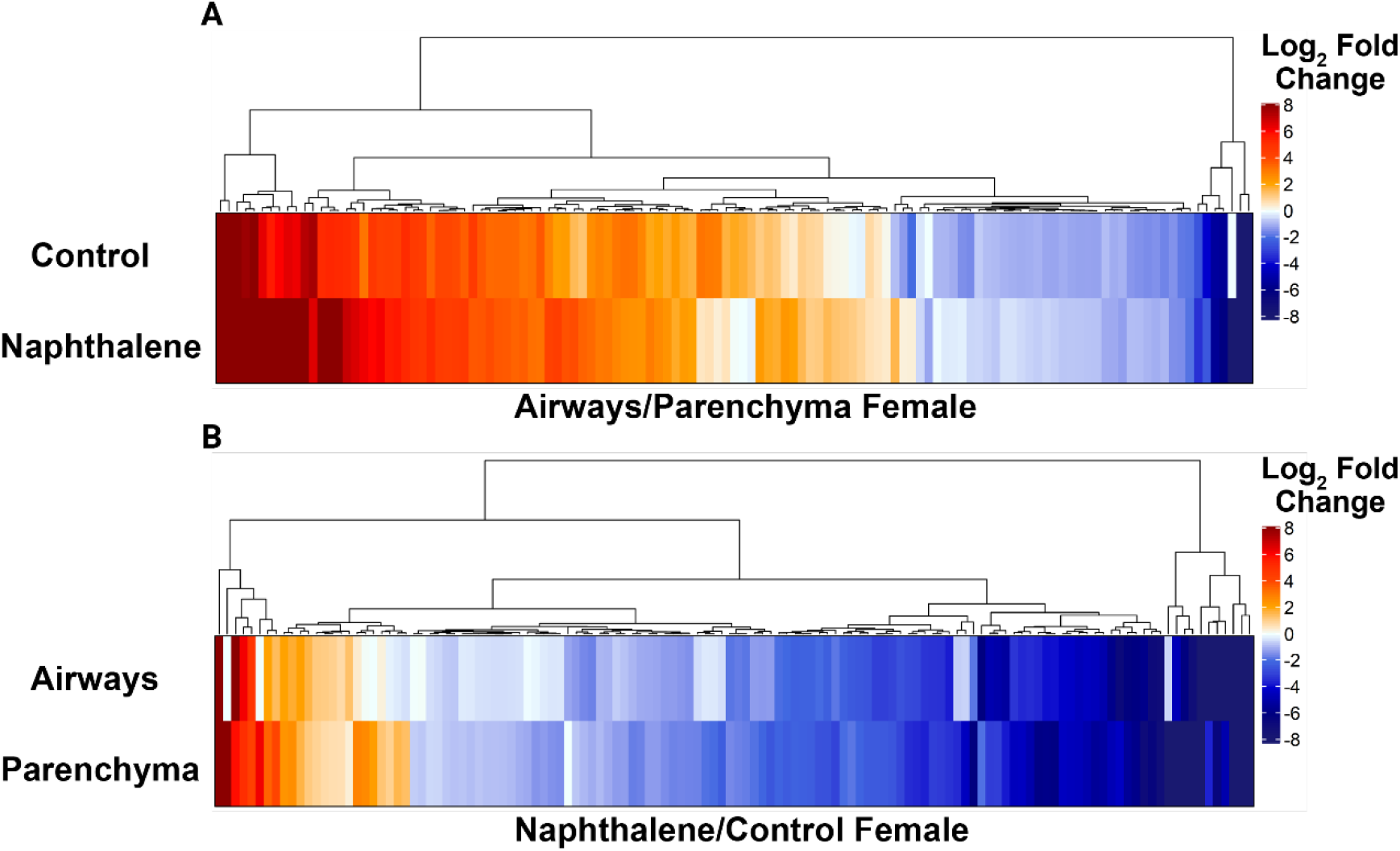
Naphthalene treatment greatly alters the profiles of individual metabolite subclasses in female airways and parenchyma at 6 hours. **A)** Heatmap comparing metabolite abundance of airways relative to parenchyma for each treatment at 6 hours. **B)** Heatmap comparing metabolite abundance of naphthalene-treated tissues relative to control-treated tissues at 6 hours for each tissue type. For both heatmaps, Euclidean clustering was used for HCA. Fold changes are expressed as the log_2_ fold change of each metabolite to indicate direction. Only metabolites that were statistically significant in at least one comparison were included in each figure. P-values were calculated by one-way ANOVA and Tukey’s post-hoc analysis. Lists of metabolites present in each heatmap are included in **Table S3** and **Table S4**.

HCA was also performed for clustering of significant changes comparing the effects of naphthalene treatment on female airways and parenchyma sampled 6 hours post-injection (**Figure 4B**). Purine and pyrimidine derivatives were clustered together and increased in both tissue types following naphthalene treatment, with airways experiencing greater relative increases in several species than parenchyma. Lysine-containing dipeptides and LPCs were also clustered together, which were ubiquitously decreased in response to naphthalene treatment (**Figure 4B**). The full lists of significantly altered metabolites are included in **Table E3** and **Table E4**.

### Univariate analysis of individual metabolites significantly affected by naphthalene treatment

Lastly, we analyzed changes in individual metabolite abundance within each tissue type following naphthalene treatment to further distinguish the response of airways compared to parenchyma. For individual metabolite analysis, we also focused on female tissues sampled 6 hours post-injection as this timepoint included the greatest number of significantly altered metabolites between sexes and each timepoint. In both airways and parenchyma, adenosine 5’-diphosphoribose, riboflavin, cytidine 5’-diphosphate ethanolamine, and uridine diphosphate galactose were all altered following naphthalene treatment, passing a threshold log_2_ fold change of 5 (**Figure 5 A-B**). However, the fold change of cytidine 5’-diphosphate ethanolamine, adenosine 5’-diphosphoribose, and uridine diphosphate galactose were all relatively greater in airways than in parenchyma, further indicating a tissue-specific response to treatment. Moreover, dipeptides containing lysine residues displayed greater relative fold changes in airways relative to changes in parenchyma. The magnitude of these changes coupled with their biological function may potentially contribute to some of the region-specific effects of naphthalene in mice.

**Figure 5.**
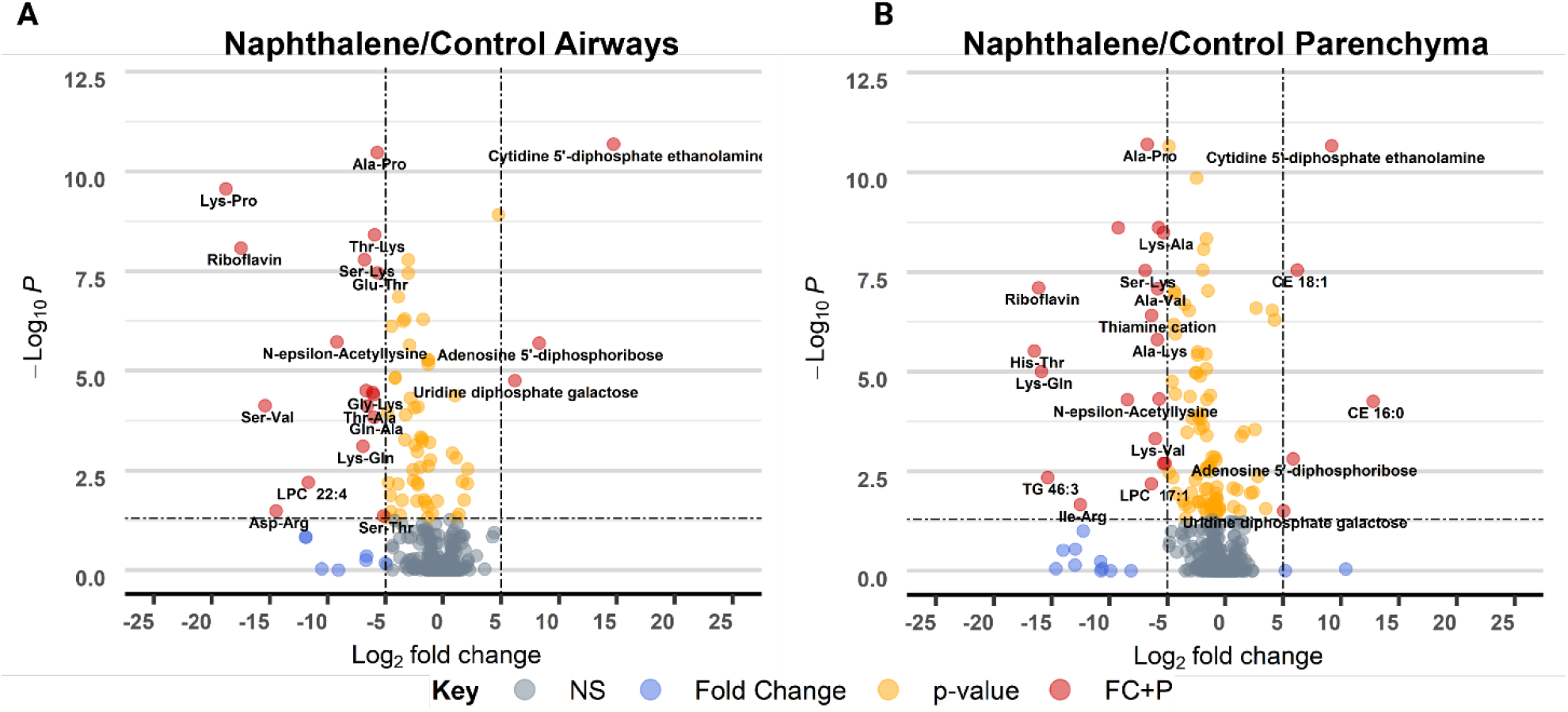
Individual metabolite changes in naphthalene-treated female mice differ in magnitude and between tissues. **A-B)** Volcano plot of −log_10_ p-value versus log_2_ fold change of metabolites in naphthalene-treated airways and parenchyma relative to control, respectively. P-values were determined using one-way ANOVA with Tukey’s post-hoc analysis. An arbitrary log_2_ fold change cutoff of 5 was used to indicate metabolites with especially large differences between treatment groups. A p-value threshold of <0.05 was selected to indicate statistical significance. Metabolites that pass both thresholds are indicated in red, whereas metabolites not passing either threshold are shaded in grey. Yellow and blue dots represent metabolites that only pass either the p-value or fold change threshold, respectively.

## Discussion

Our study demonstrates the importance of region-specific metabolomic analysis of the lung. Previous metabolomics studies of the lung have analyzed homogenized whole lung tissue to characterize the effects of naphthalene in mice (16, 17, 28). However, the results of our study demonstrated significant differences when comparing individual regions of microdissected lung tissue from male and female mice that received i.p. injections of naphthalene. PCA displayed clear separation of lung airways and parenchyma regardless of sex, treatment, or time (**Figure 2A**). Significant variation was present within each tissue, which was most likely attributable to significant differences in metabolite classes between treatments and timepoints sampled within the study.

ChemRICH analysis and HCA identified metabolite classes and subclass abundances that were unique based on tissue (**Figure 3, Figure 4**). Unsaturated TGs, PCs, and CEs were the predominant metabolite classes that varied in abundance between lung airways and parenchyma, with the relative abundance of unsaturated TGs being much greater in airways compared to PCs and CEs that were less abundant in airways. The relative abundance of these classes following naphthalene treatment shifted significantly, as unsaturated TGs greatly increased in abundance and differences between PCs and CEs became less marked in females at 6 hours (**Figure 4A**). These changes may reflect remodeling of the epithelial cell membrane and lipolysis following cytotoxicity and damage to the epithelium resulting from naphthalene treatment (5, 29). Furthermore, intake and export of TGs is dependent upon activity of apolipoprotein E (Apo-E) and apolipoprotein A-I (Apo-AI), respectively. Both proteins are expressed in the lung and serve important roles in maintaining normal lipid metabolism (30). Alterations in either Apo-E or Apo-AI are associated with several lung diseases and contribute to increased lung inflammation, oxidative stress, and collagen deposition (31–33). Although these studies have not examined the effect of naphthalene treatment on Apo-E and Apo-AI expression, selective TG accumulation in the airways of naphthalene-treated mice may suggest dysregulation of one of these proteins.

Metabolite classes and subclasses were also significantly altered when comparing the effects of naphthalene treatment in females at 6 hours post-injection. LPCs and dipeptides were the predominant classes affected by treatment, with dipeptides containing lysine residues constituting several significant differences reported in dipeptide abundance (**Figure 4B**). LPCs are bioactive lipids formed from phospholipase A_2_, which can modulate inflammatory responses and are implicated in lung disease (34, 35). LPCs can undergo conversion to phosphatidylcholines through lysophosphatidylcholine acyltransferase (LPCAT) or can be modified by another enzyme, autotaxin, to lysophosphatidic acid (LPA) (36). Previous studies have established a potential role of LPA in the development of pulmonary fibrosis through pharmacologic inhibition of the LPA receptor 1, which reduced disease severity in a bleomycin mouse model of pulmonary fibrosis (37, 38). Significant reductions in LPC following naphthalene treatment may result from increased LPA production or could result from increased production of phosphatidylcholines from LPCs through the Lands’ Cycle (**Figure 6A**) (34). Concurrent decreases in many glycine, lysine and proline-containing dipeptides may further support lung remodeling in response to naphthalene treatment considering the role of lysine and proline as important constituents of the extracellular matrix within the lung (39).

**Figure 6.**
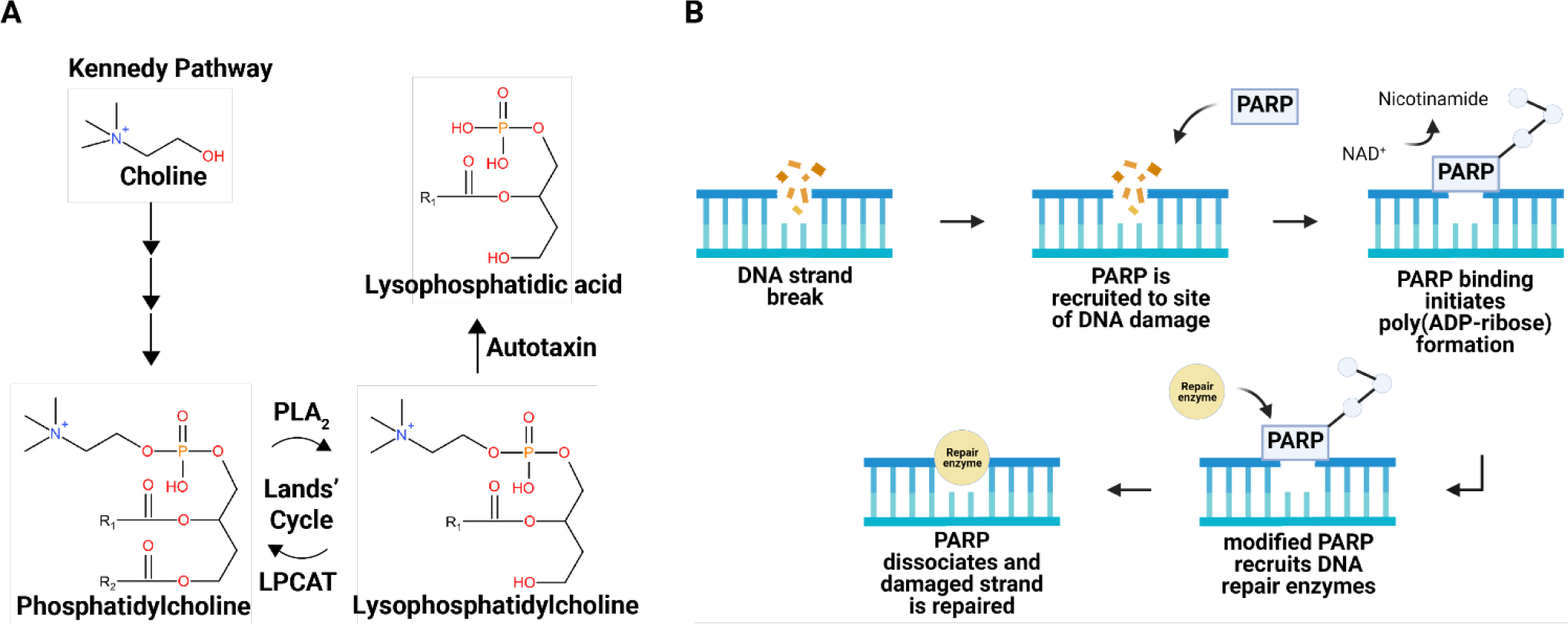
Naphthalene alters metabolites related to lung remodeling, oxidative stress, and DNA damage. **A)** De novo synthesis of phosphatidylcholine via the Kennedy pathway and subsequent breakdown into lysophosphatidylcholine by phospholipase A_2_ (PLA_2_). Lysophosphatidylcholine can either be metabolized by autotaxin into lysophosphatidic acid or converted back into phosphatidylcholine through lysophosphatidylcholine acyltransferase (LPCAT). **B)** ADP-ribose is a subunit of poly(ADP-ribose), which is formed by poly adenosine diphosphate ribose polymerase (PARP) in response to DNA strand breaks. Created with Biorender.com.

Comparisons of individual metabolites were drawn to provide insights into metabolites with unique biological functions in addition to metabolite class changes. Univariate analysis displayed striking alterations in many amino acids and pyrimidine derivatives in both female naphthalene-treated airways and parenchyma at 6 hours (**Figure 5 A-B**). Among the metabolites most substantially altered were uridine diphosphate galactose, cytidine 5’-diphosphate ethanolamine, and adenosine 5’-diphosphoribose. Interestingly, the magnitude of fold changes for each of these metabolites was much greater in airways relative to the fold change between naphthalene and control-treated parenchyma. Cytidine 5’-diphosphate ethanolamine is an important precursor used in the synthesis of phosphatidylethanolamines, which are essential components of the cell membrane (40). The significant increase of this compound in naphthalene treated mice recapitulates the observed changes in other metabolites following treatment, further implicating remodeling of the cell membrane as an effect of treatment.

Changes in cell membrane characteristics and remodeling may be exacerbated by the effects of significantly reduced lysine and significantly increased adenosine 5’-diphosphoribose also observed following treatment. Lysine has previously been reported to reduce the severity of lipopolysaccharide-induced acute lung injury by reducing lipid peroxidation and proinflammatory responses in mice (41). Reduced abundance of lysine resulting from naphthalene treatment may contribute to increased oxidative stress and glutathione depletion observed in previous studies (12). In addition to glutathione depletion, naphthalene metabolites can form DNA adducts that lead to further cytotoxicity (6). The increased abundance of adenosine 5’-diphosphoribose, which is a subunit of poly(ADP-ribose) upon DNA damage by poly adenosine diphosphate ribose polymerase (PARP), provides additional evidence supporting these previous findings (**Figure 6B**) (42–44).

Significant metabolite changes within each tissue following naphthalene treatment were almost exclusively confined to earlier timepoints, with few significant differences remaining between treatment groups 24 hours post-injection. This observation may allude to compensatory mechanisms present soon after naphthalene treatment when cells undergo vacuolization and become permeable, which contrasts with the 24-hour timepoint where club cells have exfoliated from the epithelial membrane and many are no longer present (45). Strikingly, significant metabolic changes were not limited to the airways of naphthalene-treated mice, with similar alterations being present in both airways and parenchyma (**Figure 3**). Club cells are the primary target for naphthalene toxicity in the mouse lung due to relatively high expression of CYP450 isoforms catalyzing the formation of reactive naphthalene metabolites (11). However, cell-to-cell communication is essential for maintaining homeostasis in the lung in response to injury and could potentially affect metabolic responses of other cell types such as those found in the lung parenchyma (46).

When comparing metabolite changes between each sex, male mice displayed significant alterations in multiple metabolite classes at 2 hours post-injection (**Figure E2**). In both males and females, the number of significantly altered metabolites was much lower 24 hours post-injection (data not shown). However, sex-specific differences in metabolism were most evident when comparing the effect of treatment in lung airways and parenchyma across timepoints. Changes in pyrimidine nucleotide sugars, LPCs, and dipeptides were limited to the 2 hours post-injection in males, whereas these metabolites were significantly altered at both 2 hours and 6 hours in females (**Figure E3, Figure 3**). Significant changes in metabolic profiles of male and female mice were mostly returned to baseline at 24 hours post-injection in both tissues when comparing treatments. It is well established that susceptibility to naphthalene toxicity is greater in female mice than in male mice (13, 45). These observations may be attributed to the persistence of metabolic changes related to DNA damage, oxidative stress, and lung remodeling in females but not in males. Furthermore, metabolite changes in males may also indicate protective mechanisms that mitigate naphthalene toxicity compared to females, although this was not a primary focus of our analysis.

The acute toxic effects of naphthalene exposure are well characterized in mice. However, few studies have utilized metabolomics to evaluate the contribution of global lung metabolite changes underlying the mechanism of naphthalene toxicity (7, 16, 28). Lung metabolomics studies in mice routinely analyze homogenized whole lung lobes, which prevents the characterization of region-specific responses following exposure to toxicants that target individual regions of the lung. The objective of our study was to identify region-specific differences in metabolite profiles from microdissected lung airways and parenchyma of naphthalene treated mice. The findings from this study identified inherent differences between the metabolite profiles of lung airways and parenchyma, which were further altered by naphthalene treatment. We also found significant differences in multiple metabolite classes related to oxidative stress, DNA damage, and membrane remodeling in both airways and parenchyma treated with naphthalene. Importantly, the responses between male and female mice varied greatly with respect to the duration and extent of significant changes in metabolite profiles, further validating the findings of previous studies. Future experiments are needed to examine the effects of chronic naphthalene exposure to determine the persistence of acute metabolite changes that could influence the development of lung disease. Nonetheless, the characterization of differences in lung metabolite profiles between lung airways and parenchyma in our study underscores the importance of region-specific metabolomic analysis of lung responses to target-specific toxicants.

## Supporting information

supplemental datasheet

## Acknowledgements

We would like to thank the members of the Van Winkle lab for their assistance with the execution of the animal protocol in this study. We also thank the members of the UC Davis Air Pollution and Lung Biology Journal Club for their review and suggestions provided for this manuscript.

## Supplementary Material

### Materials and Methods

#### Animal protocol

Adult male and female C57Bl6 mice (Envigo, Inc.) aged 8-10 weeks were housed on a 12/12 light/dark cycle and fed a diet consisting of Purina 5001 lab diet. Each animal received i.p. injections of either corn oil, which was used as a vehicle control, or naphthalene dissolved in corn oil (200 mg/kg) at the same time each morning. Mice were euthanized at 2, 6, or 24 hours post-injection with a lethal injection of pentobarbital and necropsied immediately following euthanasia. Lungs from each mouse were cannulated, removed en bloc, and inflated using a heated solution of 1% agarose (w/v) in 0.01M phosphate buffered saline (PBS). The left lobe of each mouse was microdissected following a previous protocol (19). The resulting airways and parenchyma were immediately stored at −80°C until analysis. All animal experiments were conducted under approved protocols reviewed by the UC Davis Institutional Animal Care and Use Committee in accordance with guidelines for animal research established by the National Institutes of Health.

#### Preparation of samples for LC/MS/MS analysis

Frozen airways and parenchyma sections were lyophilized using a Labconco freeze dryer for approximately 24 hours. Dried samples were homogenized using a mechanical disrupter (Geno/Grinder®), and 1 mg of dried tissue was utilized for analysis, which was roughly equivalent to 10 mg of fresh tissue. Tissue homogenates were extracted on ice in 2mL centrifuge tubes by adding 225 μL of methanol and an internal standard mixture included in **Table E1** and 750 μL of methyl tert-butyl ether containing cholesterol ester 22:1 (20). Samples were vortexed for 10 seconds and mixed using an orbital mixer for 5 minutes at 4°C. Each tube received 188 μL of LC-MS grade water, and the samples were vortexed for an additional 20 seconds and centrifuged at 14,000 rcf for 2 minutes. The upper hydrophobic phase was transferred into 1.5-mL centrifuge tubes each containing 350 μL for lipidomic analysis. The bottom aqueous phase was transferred into 1.5-mL centrifuge tubes for analysis of hydrophilic metabolites. A portion of the remaining upper hydrophobic phase (75 μL) from each airway and parenchyma sample was pooled into a single centrifuge tube and vortexed for 20 seconds. Individual pooled sample tubes were prepared by adding 350 μL of the pooled volume and were used as quality controls in the analysis. All samples were evaporated to dryness using a Labconco CentriVap.

The dried samples containing lipids were resuspended in 100 μL of 90% methanol, 10% toluene, and 50 ng/mL 12-[[(cyclohexylamino)carbonyl]amino]-dodecanoic acid (CUDA). The dry samples containing hydrophilic metabolites were resuspended in 110 μL of 80% acetonitrile, 20% water, and an internal standard mixture included in **Table E2**. The samples were vortexed for 10 seconds, sonicated at room temperature for 5 minutes, and centrifuged at 16,000 rcf for 2 minutes. The resuspended samples were pooled by transferring 10 μL from each airway and parenchyma sample into a single centrifuge tube, followed by mixing for an additional 10 seconds. Vials were prepared by transferring 90 μL of the resuspended samples from the sample tubes.

#### Iterative Exclusions

Spectra were acquired for each experimental sample for all platforms and combined using IE-Omics (21). Briefly, a single pooled sample was analyzed in each platform, and an R-script was used to select precursors based on ddMS^2^ topN analysis. The script was also used to generate an exclusion list for subsequent injections of the same sample. The list was imported into the instrument method to exclude the most abundant precursor ions from fragmentation in the reinjected sample. In total, 5 consecutive sample injections were run for all platforms.

#### LC/MS/MS analysis for lipids and hydrophilic metabolites

All samples were analyzed by a ThermoFisher Scientific Vanquish UHPLC+ liquid chromatography system coupled to a Q-Exactive HF orbital ion trap mass spectrometer. A Waters Acquity UPLC CSH C18 column equipped with a CSH C18 VanGuard Pre-column was used to separate metabolites for lipidomics analysis (20). Hydrophilic metabolites were analyzed through hydrophilic interaction liquid chromatography (HILIC) using a Waters UPLC BEH Amide column, equipped with a BEH Amide VanGuard Pre-column. The sample order consisted of repeated injections of one extraction blank, one bioreclaimed plasma sample, one pool of each experimental sample, followed by 10 experimental samples that were randomized prior to acquisition. For both lipidomics and HILIC analysis, 3 μL sample was injected for positive electrospray ionization and 6 μL sample for negative electrospray ionization mode. Mobile phase composition for lipid analysis consisted of 60/40 v/v acetonitrile:water (A) and 90/10 v/v isopropanol:acetonitrile (B). Modifiers used for positive mode were 0.1% formic acid and 10 mM ammonium formate, and 10mM ammonium acetate was the only modifier used for negative mode. A nonlinear gradient was run for 14.2 minutes at a constant flow rate of 0.6 mL/min. starting at 15% B 0-2 minutes, 30% B 2-2.5 minutes, 48% B 2.5-11 minutes, 82% B 11-11.5 minutes, 99% 11.5-12 minutes, 99% B 12-12.1 minutes, 15% B 12.1-14.2 minutes. The data acquisition parameters for positive mode were: 65°C column chamber temperature, 65°C post-column cooler temperature, 65°C column preheater temperature, 13-minute acquisition time, 120-1700 mass-to-charge (m/z) acquisition mass range, and stepped normalized collision energies of 20, 30, and 40%. The parameters for negative mode were identical except the acquisition mass range, which was from 113.4 to 1700 m/z.

Mobile phase composition for HILIC analysis consisted of 100% water (A) and 95/5 v/v acetonitrile:water (B) with 10mM ammonium formate and 0.125% formic acid for both ionization modes. A nonlinear gradient was run for 17 minutes at a constant flow rate of 0.4 mL/min. starting at 100% B 0-7.7 minutes, 70% B 7.7-9.5 minutes, 40% B 9.5-10.25 minutes, 30% B 10.25-12.75 minutes, and 100% B 12.75-17 minutes. The data acquisition parameters for both positive and negative ionization modes were: 45°C column chamber temperature, 45°C post-column cooler temperature, 45°C pre-heater temperature, 15-minute acquisition time, 60 to 900 m/z acquisition mass range, and stepped normalized collision energies of 20, 30, and 40%.

#### Data processing and Statistics

Deconvolution, peak picking, alignment, and identification for both lipidomics and HILIC data was completed in MS-DIAL v.4.18 (22). Identification for all compounds was based on mass spectra from *in silico* libraries, MassBank of North America (https://massbank.us), and NIST20. Experimental spectra from MassBank of North America are publicly available and NIST20 spectra are commercially available for use. Matches were determined based on m/z, retention time, and MS/MS fragmentation pattern (23). The processed data were normalized using Systematic Error Removal in Random Forest (SERRF) (24), a machine-learning algorithm that normalizes experimental samples based on systematic variation in pooled QC samples. Statistical analysis was completed in R v.3.6.1 on the log-transformed peak heights of each compound. One-way ANOVA was performed with Tukey’s HSD post-hoc test to adjust for multiple comparisons. Multivariate statistical analysis was conducted by principal component analysis (PCA), hierarchical clustering analysis (HCA) and chemical similarity enrichment analysis (ChemRICH) (25). Volcano plots and heatmaps were generated in R using the Bioconductor packages EnhancedVolcano and ComplexHeatmap (26, 27).

**Table E1.** Internal standards used in lipidomics analysis.

| Internal Standard |
| --- |
| Cholesterol ester 22:1 |
| Cholesterol-d7 |
| CUDA |
| Diacylglycerol 12:0/12:0/0:0 |
| Diacylglycerol 18:1/2:0/0:0 |
| Lysophosphatidylcholine 17:0 |
| Lysophosphatidylethanolamine 17:1 |
| Monoacylglycerol 17:0/0:0/0:0 |
| Phosphatidylcholine 12:0/13:0 |
| Phosphatidylethanolamine 17:0/17:0 |
| Sphingomyelin d18:1/17:0 |
| Sphingosine d17:1 |
| Triacylglycerol (14:0/16:1/14:0)-d5 |
| Triacylglycerol d5 17:0/17:1/17:0 |
| 5-PAHSA-d9 |
| Ceramide d18:1/17:0 |
| Fatty acid 16:0-d3 |
| Phosphatidylglycerol 17:0/17:0 |
| Phosphatidylinositol (15:0-18:1)-d7 |
| Phosphatidylserine (15:0-18:1)-d7 |

**Table E2.**
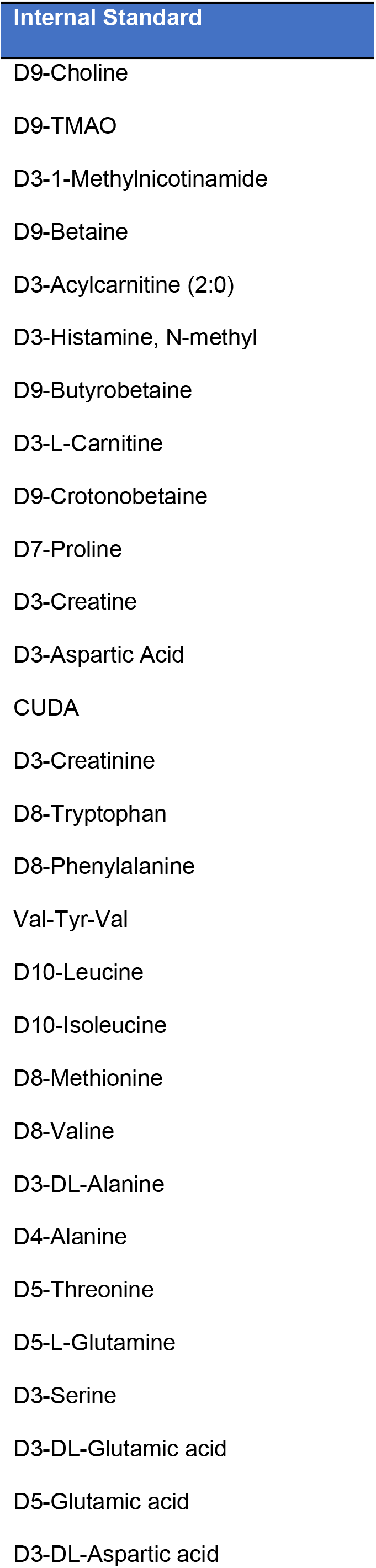
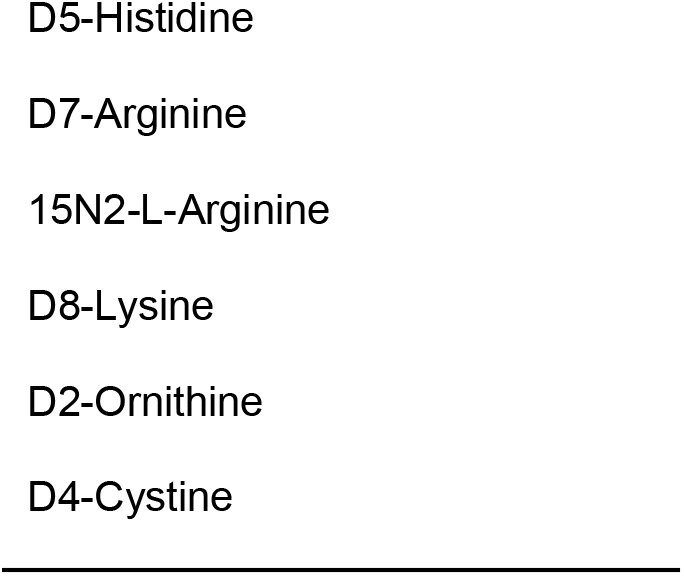
Internal standards used in HILIC analysis.

**Table E3.**
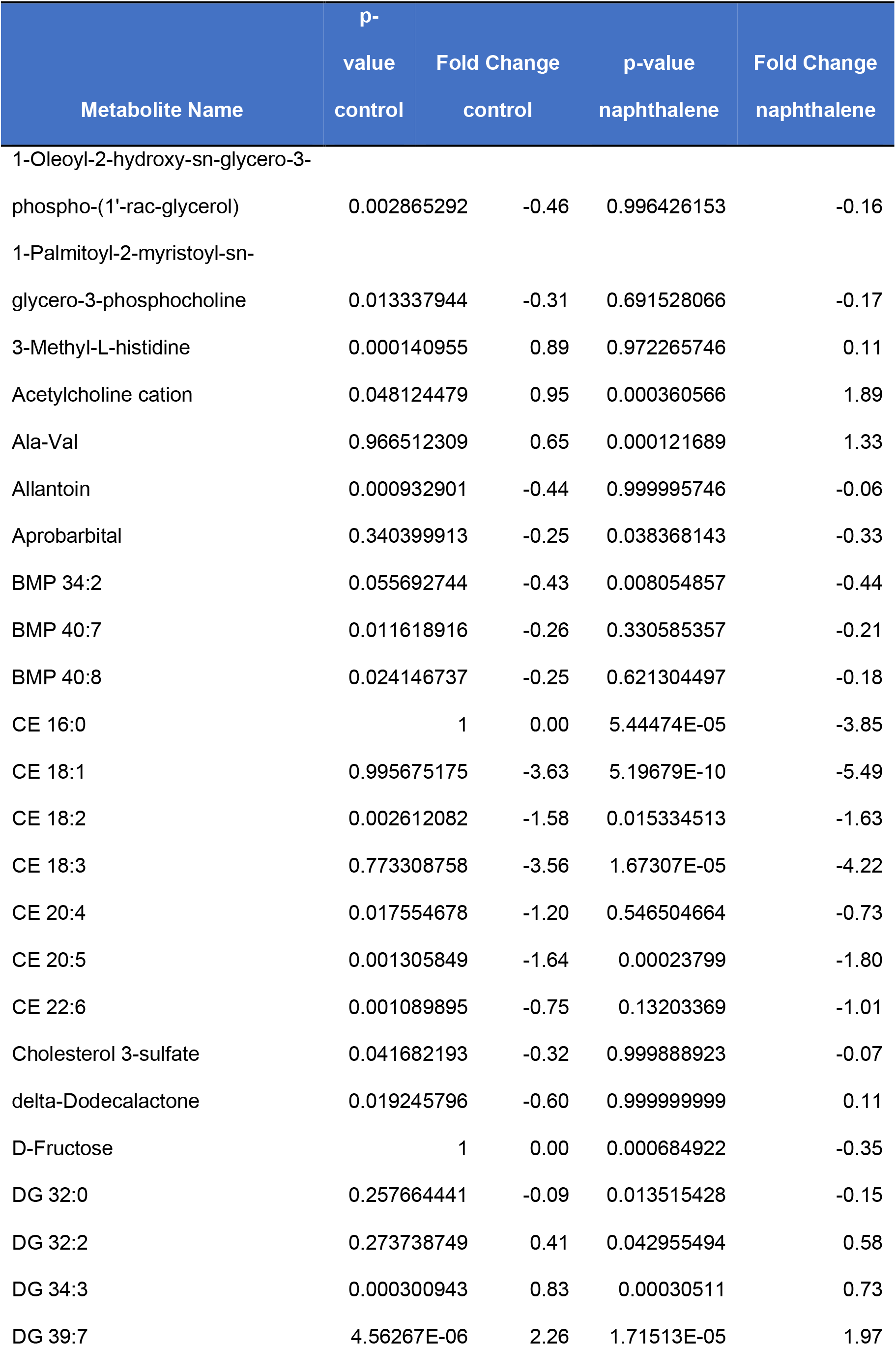
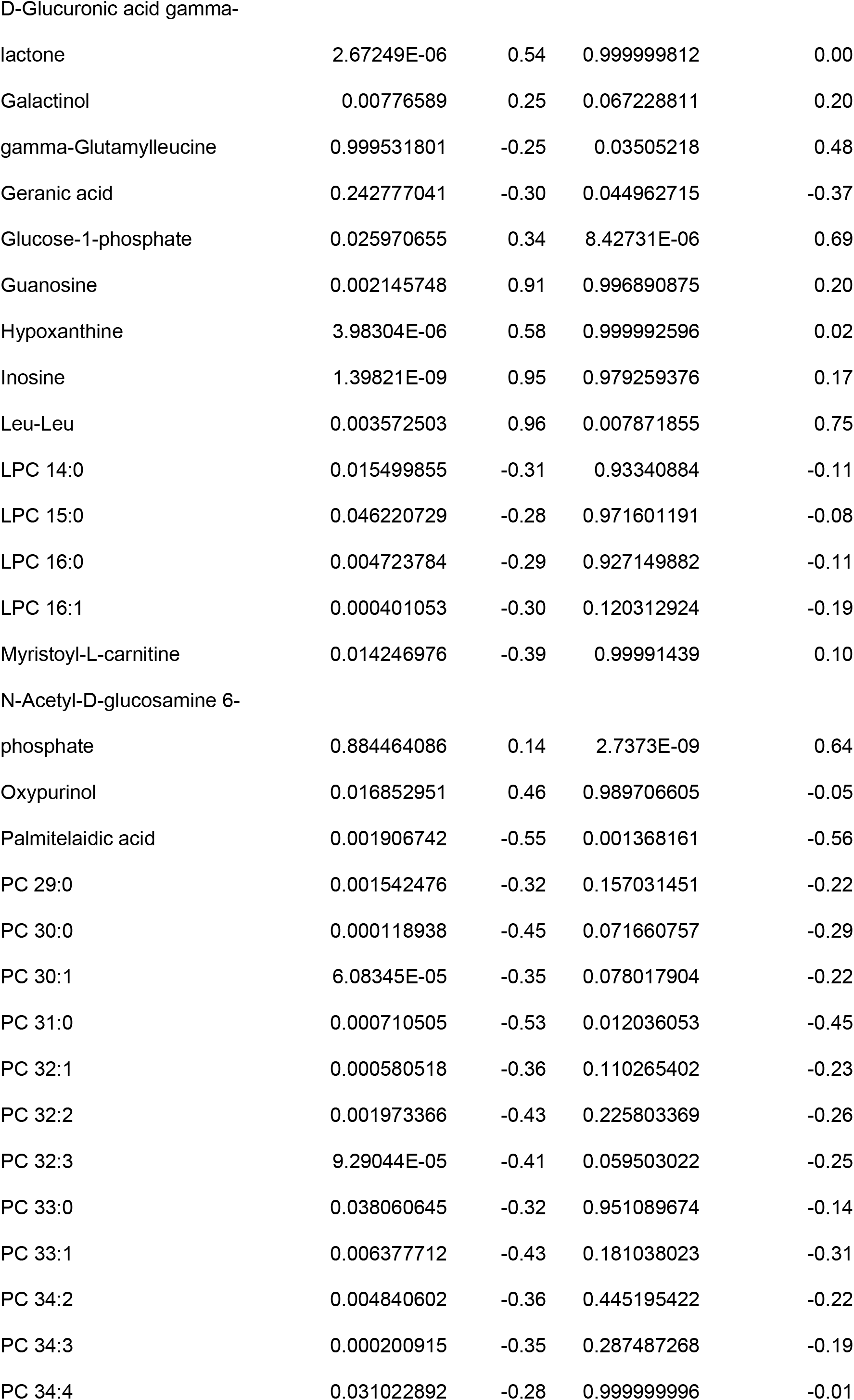
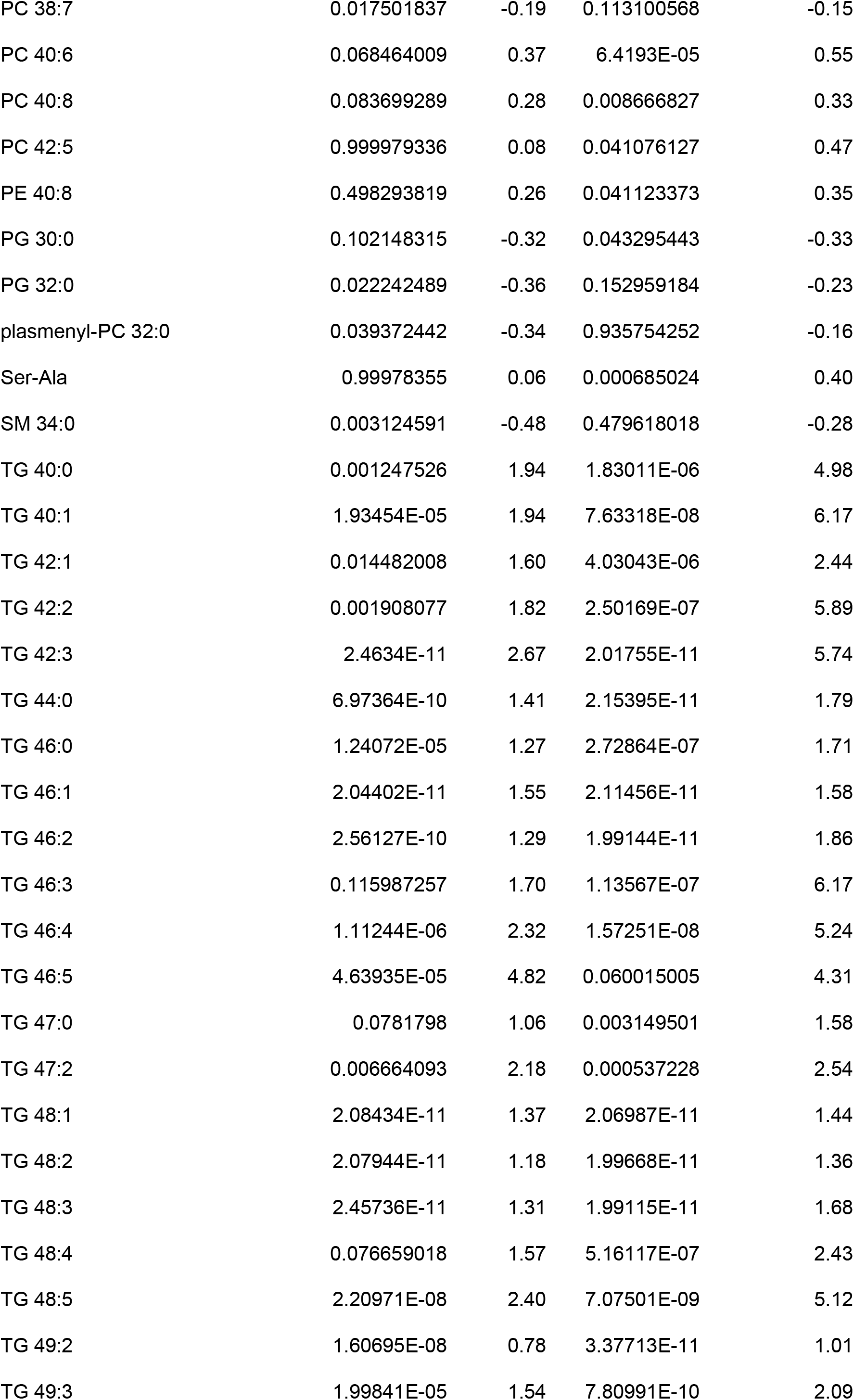
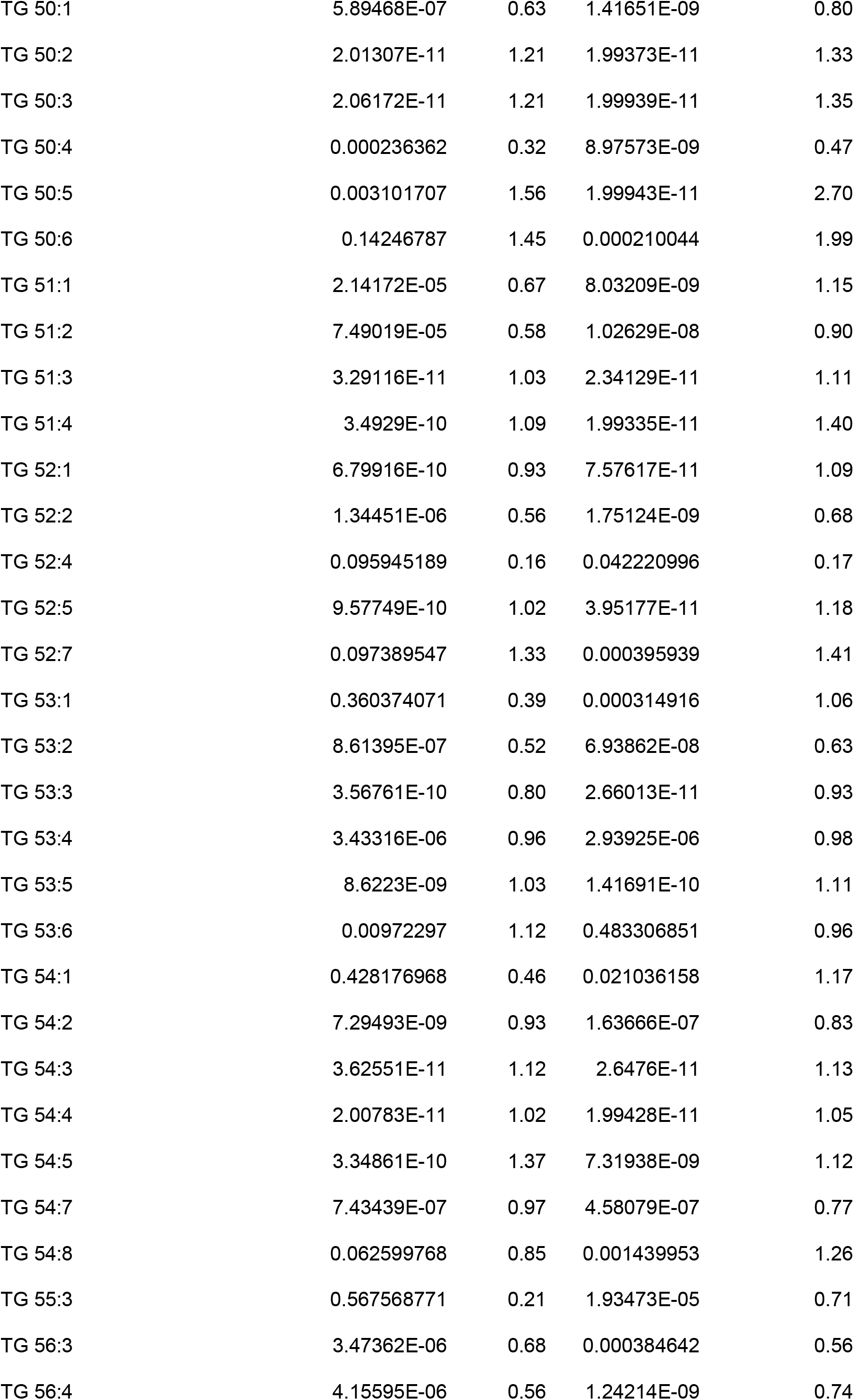
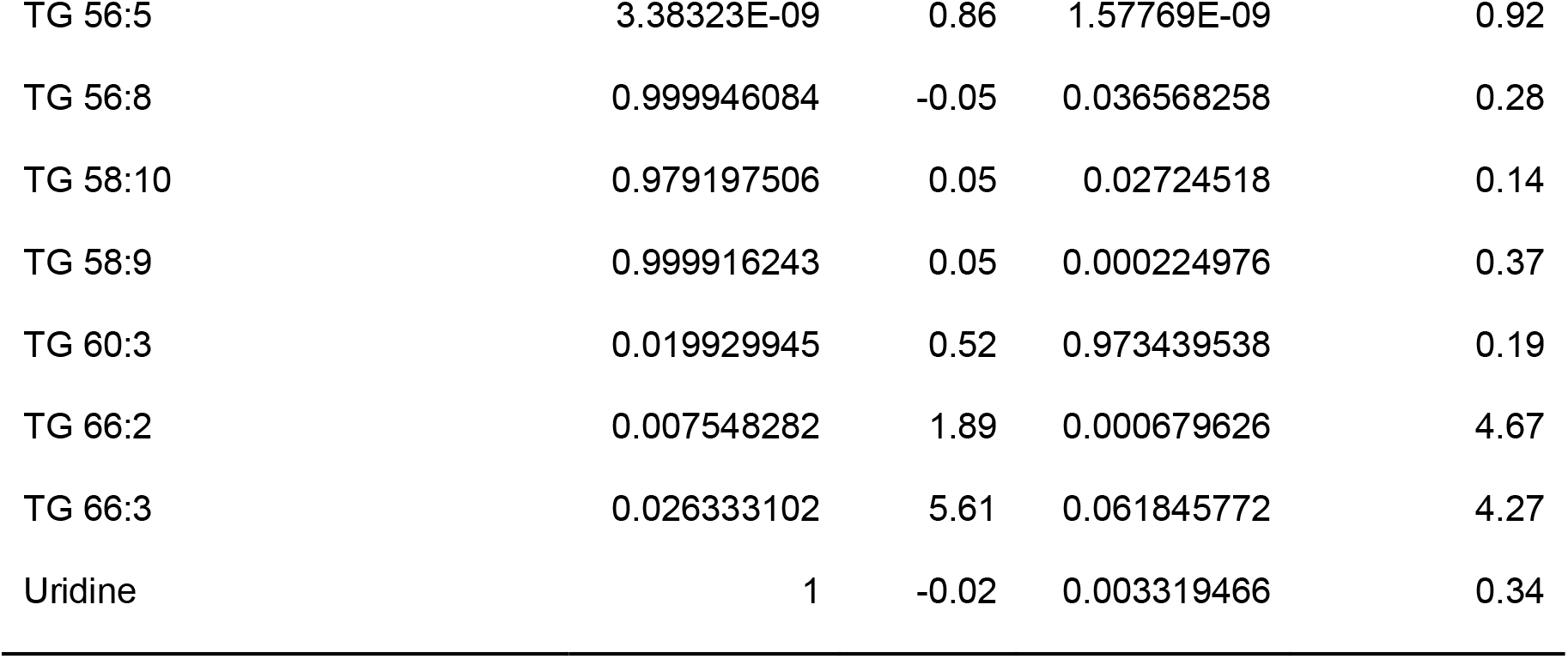
Metabolites significantly different between female mouse airways and parenchyma at 6 hours post-injection for each treatment group.

**Table E4.** Metabolites significantly different between naphthalene- and control-treated female mice at 6 hours post-injection for each tissue.

| Metabolite Name | p-value<br>airways | Fold change<br>airways | p-value<br>parenchyma | Fold Change<br>parenchyma |
| --- | --- | --- | --- | --- |
| 12-oxo-ETE | 0.883353626 | -0.31 | 0.002363285 | -1.43 |
| 13,14-Dihydro-15-ketoprostaglandin A2 | 0.02533179 | -1.35 | 5.02983E-05 | -1.59 |
| 13-Keto-9Z,11E-octadecadienoic acid | 0.090515024 | -1.44 | 0.000396236 | -1.62 |
| 14-HDoHE | 0.121186491 | -1.31 | 0.000112629 | -2.11 |
| 1-Oleoyl-2-hydroxy-sn-glycero-3-phospho-(1'-rac-glycerol) | 0.039611354 | 1.36 | 1 | 0.36 |
| 3-Methyl-L-histidine | 0.14926271 | 1.48 | 2.93419E-07 | 4.06 |
| 8-Oxo-2-deoxyadenosine | 0.017489747 | 1.81 | 0.000281334 | 2.57 |
| 9-Oxo-10(E),12(E)-octadecadienoic acid | 0.957582444 | -0.78 | 0.030457479 | -1.27 |
| Acetylenedicarboxylic acid | 0.018277584 | -0.50 | 0.025543071 | -0.51 |
| Adenosine 5'-diphosphoribose | 2.05348E-06 | 8.28 | 0.001522438 | 5.86 |
| Adenosine 5'-monophosphate | 0.639454573 | 2.09 | 0.02709777 | 3.50 |
| Ala-Ala | 0.503097621 | -2.03 | 0.018257852 | -2.36 |
| Ala-Lys | 1.40258E-07 | -3.86 | 1.58163E-06 | -5.88 |
| Alanine | 0.003025151 | -2.59 | 0.000146798 | -2.87 |
| Ala-Pro | 3.3395E-11 | -5.74 | 1.99586E-11 | -6.74 |
| Ala-Thr | 0.001062654 | -2.22 | 0.017263127 | -1.96 |
| Ala-Val | 0.069621108 | -3.58 | 8.21715E-08 | -5.84 |
| Allantoin | 1 | -0.11 | 0.003197828 | -1.39 |
| Arginine | 5.09689E-07 | -3.32 | 2.95778E-07 | -3.12 |
| Arg-Thr | 0.84548275 | -3.25 | 0.016892245 | -3.66 |
| Asp-Arg | 0.032637624 | -14.45 | 0.188557985 | -4.81 |
| Bestatin | 1.63778E-08 | -3.01 | 3.21815E-06 | -2.38 |
| beta-Homoproline | 0.006729921 | 2.08 | 0.010513068 | 2.41 |
| Bicyclo-prostaglandin E2 | 0.999894693 | -0.30 | 0.002483754 | -1.48 |
| CE 16:0 | 1 | 0.00 | 5.54094E-05 | 12.78 |
| CE 18:1 | 1 | 0.05 | 2.80136E-08 | 6.23 |
| CE 18:3 | 1 | 0.01 | 0.010996891 | 2.19 |
| Choline | 0.048190857 | -1.33 | 0.33843499 | -0.72 |
| Cystine | 0.00012837 | -3.23 | 9.58729E-08 | -4.41 |
| Cytarabine | 0.006506357 | -2.22 | 0.000270123 | -2.16 |
| Cytidine 2',3'-cyclic monophosphoric acid | 8.42805E-05 | -2.48 | 0.000154453 | -2.15 |
| Cytidine 5'-diphosphate ethanolamine | 2.0829E-11 | 14.73 | 2.16577E-11 | 9.19 |
| Cytidine-5'-monophosphate | 0.002898603 | 2.12 | 0.0826784 | 1.65 |
| D-Fructose | 0.000629415 | -1.16 | 1 | 0.01 |
| DG 30:0 | 0.400115113 | -0.94 | 3.65742E-06 | -1.70 |
| DG 31:0 | 0.126348157 | -0.71 | 0.033035123 | -0.78 |
| DG 32:1 | 0.069724226 | -0.67 | 0.014667333 | -0.71 |
| DG 33:1 | 0.969600963 | -0.42 | 0.02145158 | -0.81 |
| DG 36:1 | 0.879884221 | -0.37 | 0.007663538 | -0.73 |
| DG 38:5 | 0.77550267 | -0.38 | 0.001673593 | -0.81 |
| D-Glucuronic acid gamma-lactone | 0.997696339 | -0.39 | 0.000412155 | 1.40 |
| FA 16:1 | 0.019187031 | -1.55 | 0.209728183 | -0.91 |
| FA 18:2 | 0.017206422 | -1.64 | 0.13743199 | -1.06 |
| FA 20:2 | 0.15849512 | -1.15 | 0.022986149 | -1.29 |
| GlcCer 42:1 | 0.999991014 | -0.16 | 0.009111821 | 1.01 |
| Gln-Ala | 0.000146926 | -6.01 | 0.090746918 | -1.90 |
| Gln-Thr | 0.000501143 | -1.82 | 0.008447503 | -1.55 |
| Glu-Gly-Arg | 0.274534277 | -4.35 | 0.003388058 | -4.85 |
| Glu-Thr | 3.4537E-08 | -5.70 | 1.77168E-05 | -4.59 |
| Gly-Lys | 3.51156E-05 | -6.11 | 0.001989254 | -5.35 |
| Gly-Ser | 0.017648568 | -3.54 | 3.63364E-05 | -4.32 |
| Gly-Tyr | 0.000460656 | -1.93 | 1.28658E-05 | -2.15 |
| Guanosine | 0.999614596 | 0.45 | 0.004232542 | 2.81 |
| Guanosine-5'-monophosphate | 0.047608097 | 1.16 | 0.288064244 | 0.97 |
| His-Gln | 0.006324604 | -4.75 | 0.024165713 | -3.09 |
| His-Leu | 0.048140267 | -4.94 | 4.21509E-05 | -3.04 |
| His-Thr | 4.00206E-05 | -6.05 | 3.02748E-06 | -16.51 |
| Histidine | 0.018256264 | -2.26 | 0.005165274 | -2.60 |
| Hypoxanthine | 0.999954759 | -0.26 | 0.000324069 | 1.60 |
| Ile-Arg | 0.013341516 | -4.57 | 0.021465788 | -12.53 |
| Indole-3-carboxylic acid | 0.0005769 | -1.81 | 0.008826388 | -1.71 |
| Inosine | 1 | 0.07 | 2.58703E-07 | 2.67 |
| LPC 17:1 | 0.146956869 | -11.90 | 0.006443851 | -6.40 |
| LPC 20:0 | 0.971931845 | -0.38 | 0.028582468 | -1.04 |
| LPC 20:1 | 0.198362972 | -1.10 | 0.002538803 | -1.64 |
| LPC 22:4 | 0.00627216 | -11.67 | 0.166093405 | -3.60 |
| LPC 22:6 | 5.21716E-07 | -1.73 | 8.48566E-06 | -1.65 |
| LPC 14:0 | 0.592062487 | -0.63 | 0.002019848 | -1.29 |
| LPC 16:1 | 6.86109E-06 | -1.28 | 4.5679E-09 | -1.63 |
| LPC 18:0 | 0.997529992 | -0.27 | 0.035032057 | -0.84 |
| LPC 18:1 | 0.001695943 | -1.11 | 3.93357E-05 | -1.30 |
| LPC 18:2 | 5.4053E-06 | -1.28 | 9.20417E-08 | -1.50 |
| LPC 20:3 | 0.005601261 | -2.55 | 0.096249617 | -2.38 |
| LPC 20:4 | 5.41734E-06 | -1.28 | 8.4647E-09 | -1.87 |
| LPC 22:5 | 0.000538268 | -3.30 | 0.097483766 | -3.05 |
| Lys-Ala | 7.7306E-07 | -4.48 | 3.20427E-09 | -5.34 |
| Lys-Gln | 0.000780483 | -6.96 | 9.97485E-06 | -15.90 |
| Lysine | 1.54712E-05 | -4.19 | 6.5609E-07 | -4.44 |
| Lys-Pro | 2.73201E-10 | -18.78 | 2.43399E-09 | -9.25 |
| Lys-Val | 0.1683592 | -1.81 | 0.000477647 | -6.05 |
| Methionine | 2.29999E-06 | -2.91 | 1.06666E-05 | -2.50 |
| Myristoyl-L-carnitine | 0.99999193 | -0.27 | 0.000236862 | -1.89 |
| N-Acetyl-D-glucosamine 6-phosphate | 0.995571274 | -0.28 | 2.78961E-08 | -1.96 |
| N-alpha-(tert-Butoxycarbonyl)-L-histidine | 0.007243105 | -2.17 | 0.018113467 | -2.24 |
| N-epsilon-Acetyllysine | 1.90843E-06 | -9.20 | 5.08125E-05 | -8.47 |
| N-Methyl-L-proline | 1.46539E-05 | -4.14 | 1.14025E-06 | -4.32 |
| Ornithine | 0.072118775 | -1.80 | 0.010794639 | -2.93 |
| Palmitic acid | 0.001535523 | 1.12 | 0.133955178 | 0.78 |
| PC 25:0 | 0.999999598 | 0.31 | 0.031406369 | 1.27 |
| PC 38:6 | 0.131912639 | -0.41 | 0.028446805 | -0.49 |
| PE 40:4 | 0.800133419 | -0.42 | 0.011214684 | -0.83 |
| Phenylalanine | 3.52411E-08 | -3.02 | 3.84051E-06 | -2.39 |
| Pro-Leu | 0.041242594 | -3.72 | 0.000332358 | -3.31 |
| Proline | 4.89776E-05 | -2.82 | 2.03357E-07 | -3.53 |
| Pro-Pro | 0.000114928 | -4.81 | 0.001967272 | -5.19 |
| rac-Glycerol 3-phosphoate | 0.037214872 | -0.93 | 0.015981848 | -0.89 |
| Riboflavin | 8.32671E-09 | -17.47 | 7.76424E-08 | -16.14 |
| Ribulose 5-phosphate | 0.15035441 | -2.78 | 0.047482959 | -3.43 |
| Ser-Asn | 0.033939663 | -4.52 | 0.004504892 | -4.53 |
| Ser-Glu | 0.446668654 | -0.97 | 0.022448285 | -1.10 |
| Serine | 8.04009E-05 | -2.17 | 0.000137416 | -2.30 |
| Ser-Lys | 1.61517E-08 | -6.83 | 2.86041E-08 | -6.92 |
| Ser-Pro | 0.006875018 | -3.89 | 2.40476E-09 | -5.76 |
| Ser-Thr | 0.043487012 | -5.18 | 0.111902282 | -3.76 |
| Ser-Val | 7.49765E-05 | -15.39 | 0.097994985 | -12.28 |
| SM 32:1 | 0.794390434 | -0.40 | 0.001406002 | -0.89 |
| TG 46:2 | 0.999072872 | -0.41 | 0.003540875 | -2.30 |
| TG 46:3 | 1 | -0.50 | 0.004496577 | -15.35 |
| TG 48:3 | 0.994367639 | -0.42 | 0.033896396 | -1.64 |
| TG 48:4 | 0.999999999 | -0.62 | 0.021626578 | -3.47 |
| TG 50:4 | 0.991192038 | -0.25 | 0.016047761 | -0.75 |
| TG 50:5 | 0.999998247 | -0.56 | 1.12936E-07 | -4.36 |
| TG 51:4 | 0.99986825 | -0.25 | 0.020804687 | -1.29 |
| TG 56:7 | 1 | 0.05 | 0.034901515 | -0.71 |
| TG 56:8 | 0.999999648 | -0.03 | 0.00136058 | -1.11 |
| TG 56:9 | 0.99999997 | 0.01 | 0.026855358 | -0.63 |
| TG 60:11 | 0.001163844 | 0.80 | 0.008279034 | 0.69 |
| TG 60:12 | 4.2112E-05 | 1.03 | 0.024401089 | 0.71 |
| Thiamine cation | 3.15056E-05 | -6.69 | 3.89475E-07 | -6.37 |
| Thr-Ala | 7.64777E-05 | -6.60 | 0.011193507 | -4.28 |
| Threonine | 0.002598595 | -1.96 | 1.078E-05 | -2.58 |
| Thr-Lys | 3.8716E-09 | -5.96 | 4.84662E-05 | -5.72 |
| Uridine | 0.00244635 | -1.28 | 1.39152E-10 | -2.48 |
| Uridine 5'-monophosphate | 0.005959534 | 1.65 | 0.192569005 | 1.16 |
| Uridine diphosphate galactose | 1.76582E-05 | 6.18 | 0.030821525 | 5.05 |
| Uridine-5-diphosphoacetylglucosamine | 1.21668E-09 | 4.77 | 5.11274E-07 | 4.26 |
| Val-Thr | 0.000728176 | -2.40 | 0.026546975 | -1.81 |
| Xanthosine | 5.78725E-07 | -3.44 | 2.21857E-11 | -4.86 |

**Figure E1.**
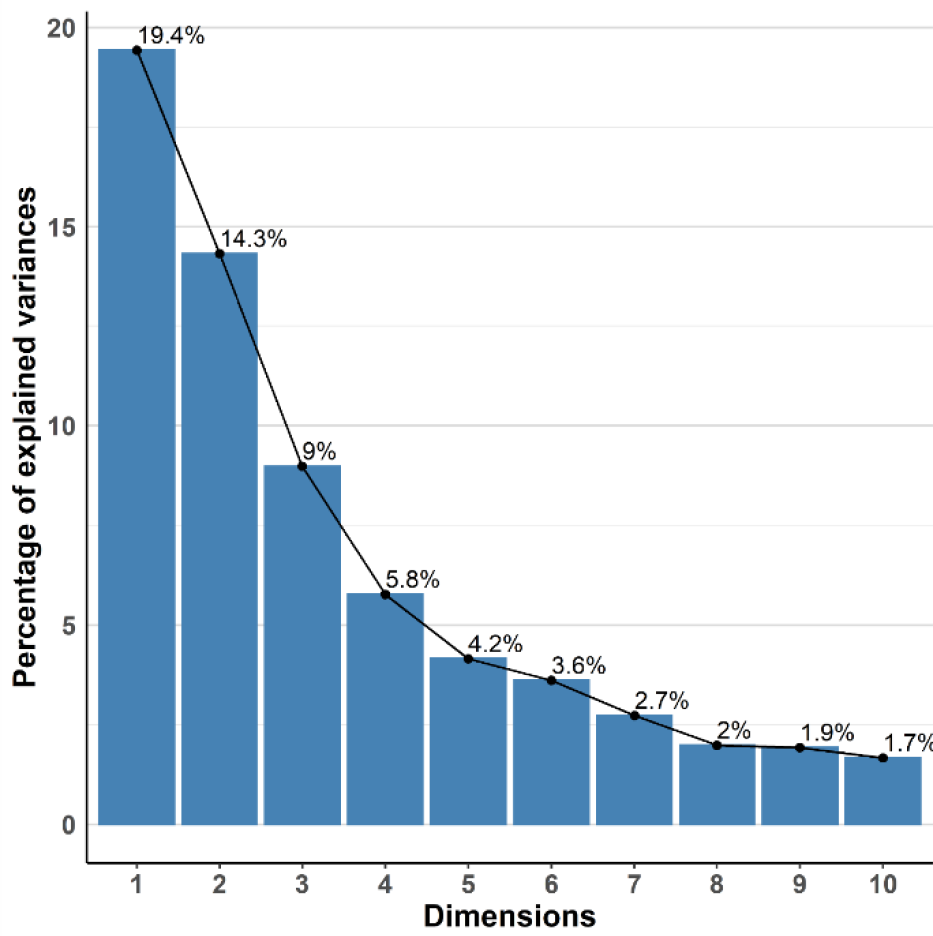
Scree plot indicating the percent of variance contained within the first 10 principal components. The percent of variance contained within the first 5 principal components is less than 50% of the total variance explained by PCA. Each bar represents a different dimension, where dimension 1 corresponds to the first principal component.

**Figure E2.**
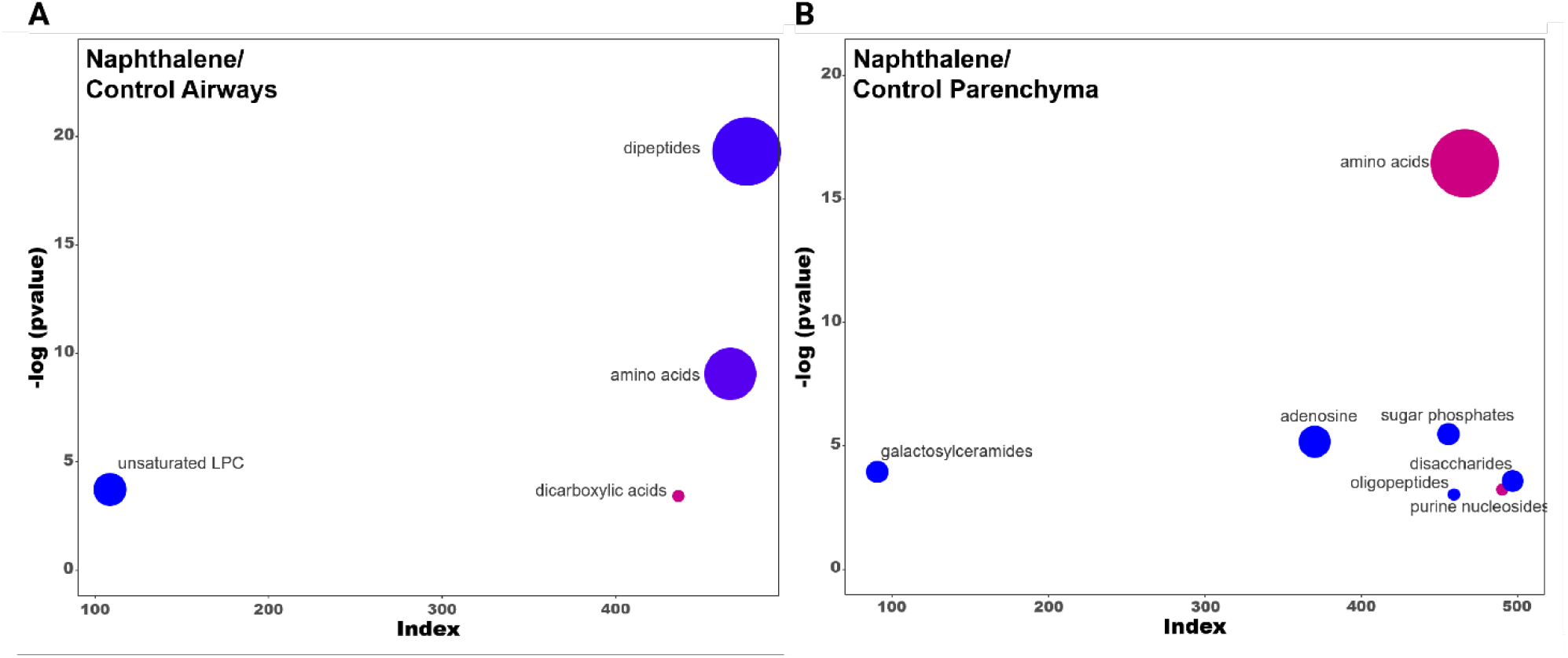
Metabolite profiles of lung airways and parenchyma are greatly altered following naphthalene treatment in males 2 hours post-injection. **A-B)** ChemRICH plots comparing naphthalene-treated airways and parenchyma in male mice 2 hours post-injection, respectively. The size of each circle represents the relative number of metabolites contained within each cluster. Red circles indicate all metabolites increase within a cluster, while blue circles indicate all metabolites decrease within a cluster. Pink and purple circles represent a mix consisting of mostly increased and decreased metabolite abundances, respectively. Axes correspond to the −logP value of a metabolite class plotted against index values assigned to each metabolite in the online datasheet included as supplemental material. P-values used for the input of each ChemRICH were calculated by one-way ANOVA with Tukey’s post-hoc analysis. P-values for each ChemRICH cluster were calculated using the Kolmogorov-Smirnov test.

**Figure E3.**
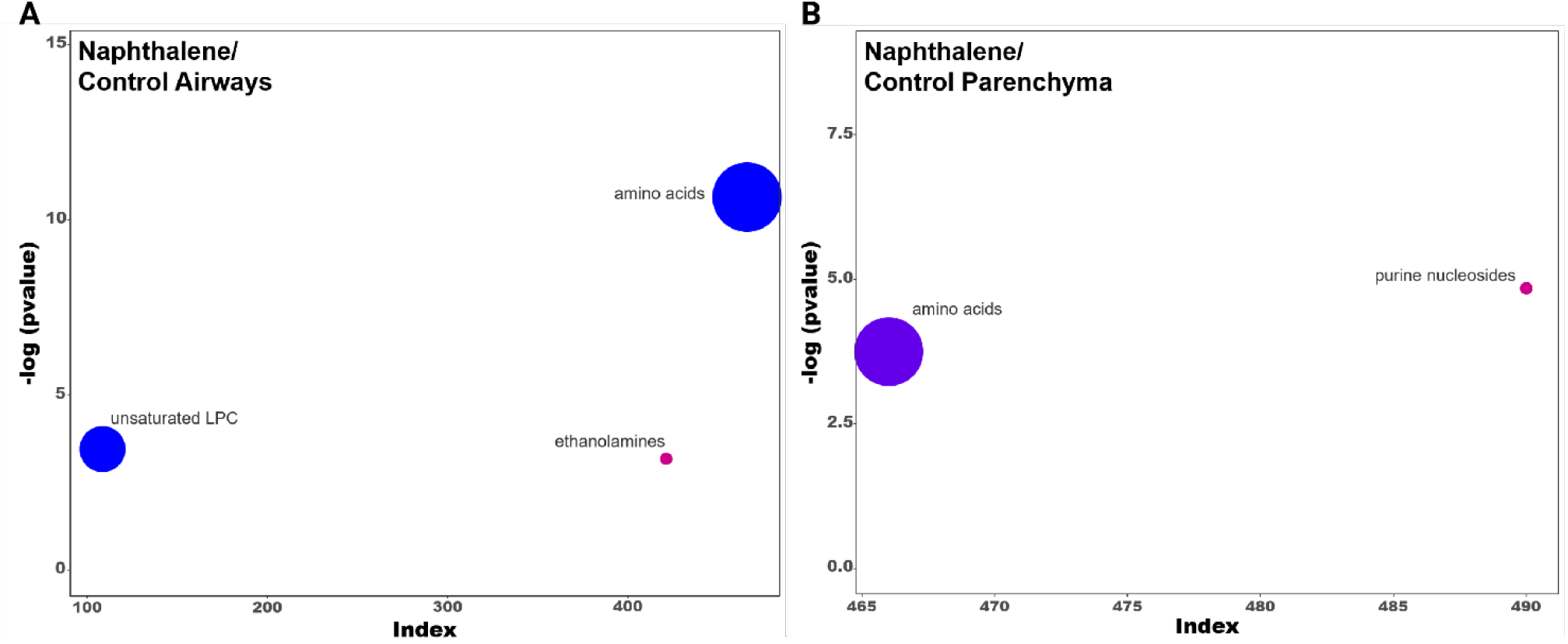
Metabolite profiles of lung airways and parenchyma are greatly altered following naphthalene treatment in females 2 hours post-injection. **A-B)** ChemRICH plots comparing naphthalene-treated airways and parenchyma in female mice 2 hours post-injection, respectively. The size of each circle represents the relative number of metabolites contained within each cluster. Red circles indicate all metabolites increase within a cluster, while blue circles indicate all metabolites decrease within a cluster. Pink and purple circles represent a mix consisting of mostly increased and decreased metabolite abundances, respectively. Axes correspond to the −logP value of a metabolite class plotted against index values assigned to each metabolite in the online datasheet included as supplemental material. P-values used for the input of each ChemRICH were calculated by one-way ANOVA with Tukey’s post-hoc analysis. P-values for each ChemRICH cluster were calculated using the Kolmogorov-Smirnov test.

**Figure E4.**
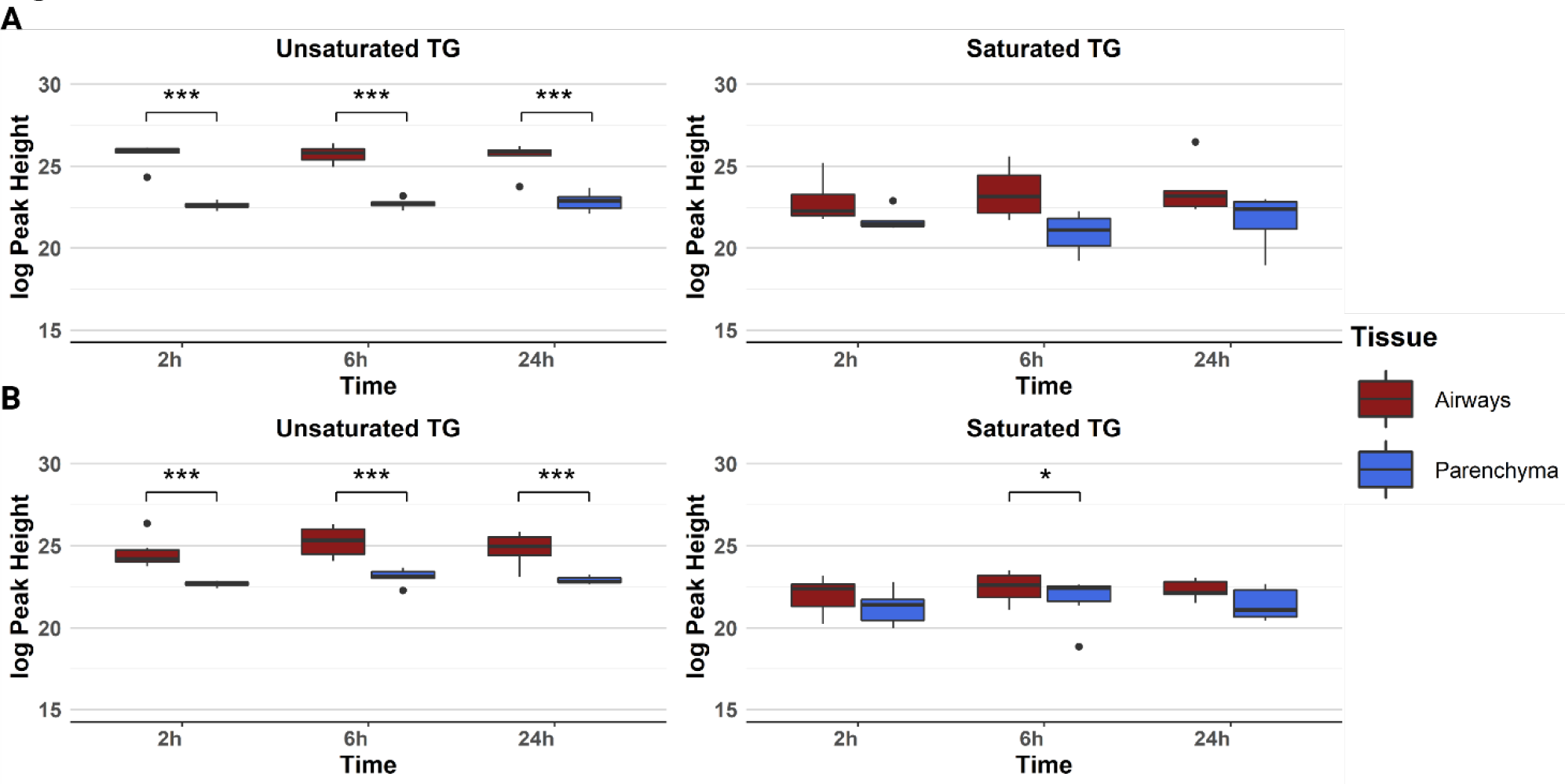
Triacylglyceride abundance does not fluctuate between different timepoints for both female and male mice. **A-B)** Boxplots displaying the average intensities for saturated and unsaturated TGs in female and male mice at each timepoint, respectively. Axes represent the log_10_ peak height of each sample for each timepoint, and samples with values greater than 1.5 times the interquartile range are indicated by dots on each plot. * p<0.05, *** p<0.001.

